# Unravelling the effect of a potentiating anti-Factor H antibody on atypical hemolytic uremic syndrome associated factor H variants

**DOI:** 10.1101/2020.04.08.026906

**Authors:** Gillian Dekkers, Mieke Brouwer, Jorn Jeremiasse, Angela Kamp, Robyn M. Biggs, Gerard van Mierlo, Scott Lauder, Suresh Katti, Taco W. Kuijpers, Theo Rispens, Ilse Jongerius

## Abstract

The complement system plays an important role in our innate immune system. Complement activation results in clearance of pathogens, immune complex and apoptotic cells. The host is protected from complement-mediated damage by several complement regulators. Factor H (FH) is the most important fluid-phase regulator of the alternative pathway of the complement system. Heterozygous mutations in FH are associated with complement-related diseases such as atypical hemolytic uremic syndrome (aHUS) and age-related macular degeneration.

We recently described an agonistic anti-FH monoclonal antibody that can potentiate the regulatory function of FH. This antibody could serve as a potential new drug for aHUS patients and alternative to C5 blockade by Eculizumab. However, it is unclear whether this antibody can potentiate FH mutant variants in addition to wild type FH. Here, the functionality and potential of the agonistic antibody in the context of pathogenic aHUS-related FH mutant proteins was investigated. The binding affinity of recombinant WT FH, and the FH variants, W1183L, V1197A, R1210C, and G1194D to C3b was increased upon addition of the potentiating antibody and similarly, the decay accelerating activity of all mutants is increased. The potentiating anti-FH antibody is able to restore the surface regulatory function of most of the tested FH mutants to WT FH levels. In conclusion, our potentiating anti-FH is broadly active and able to enhance both WT FH function as well as most aHUS-associated FH variants tested in this study.

## Introduction

The complement system plays an important role in our immune system ensuring pathogens, immune complexes and dying cells are efficiently cleared from circulation. Activation of the complement system can occur via three pathways: the classical (CP), lectin (LP) and alternative pathway (AP). Activation of any of these pathways leads to the cleavage of the central complement component C3, into C3a and C3b. C3a is an anaphylatoxin and C3b attaches covalently to surfaces, creating an opsonization signal that allows recognition and clearance by the immune system. In addition, C3b can lead to the formation of C5 convertases that cleave C5 into C5a and C5b, which is necessary to form the membrane attack complex (MAC), resulting in lysis of the target (1–3).

The AP can be initiated via the spontaneous hydrolyzation of C3 (tick-over) in the fluid phase, forming C3(H_2_O) and serves to amplify the effects of activated CP and the LP. The AP amplification loop starts with the binding of Factor B (FB) to C3b and is cleaved by Factor D (FD), forming the AP C3 convertase C3bBb, which can cleave another C3. Binding of an additional C3b molecule to the C3bBb forms the AP C5 convertase (4, 5). To prevent unwanted damage to the body’s own cells by this amplification loop, complement is tightly regulated by fluid-phase and membrane bound complement regulators (1, 2, 6). Factor H (FH) is the main regulator of the AP (7).

FH regulates C3 convertase activity both in fluid phase as well as on the cellular surface. FH binds to C3b and mammalian cellular surfaces by interaction with host specific glycosaminoglycans (GAGs) or sialic acid containing glycans on the cellular membrane (8). This interaction results in decay of the AP convertase C3bBb. Furthermore, FH together with factor I (FI) degrades C3b to iC3b and renders it proteolytically inactive towards formation of the convertases (9). FH is composed of 20 complement control protein (CCP) domains and can be further divided into regions that specifically interact with sialic acids and GAGs on the surface of human cells, C3b or both (10, 11). The N-terminal region, composed of CCP domains 1 to 4, is involved in binding to C3b and responsible for the decay accelerating activity (DAA) and co-factor activity of FH. There are additional C3b binding sites within the regions encompassed by CCPs 7 to 15 and 19 to 20, of which the latter interacts with C3b, iC3b and C3d. The FH CCPs 7, 20 and the 15-19 region are involved in heparin and polyanion binding, and the C-terminal CCP-20 is involved in sialic acid binding. The binding of FH CCPs 19 and 20 to both C3b and polyanions is crucial for the host recognition (7, 11, 12).

Atypicial hemolytic uremic syndrome (aHUS) is a complement-mediated disease, characterized by hemolytic anemia, thrombocytopenia and complement depositions in the kidney in particular, resulting in acute renal failure (13). Many aHUS patients carry genetic mutations in a complement protein gene, where 25-30% of the cases are observed to have FH gene mutations and ∼5% of aHUS cases are observed to develop blocking autoantibodies against FH (13). The aHUS patients carrying a FH mutation are heterozygously affected (14, 15). Whether both WT and mutant variant of FH are equally expressed and thus present in the patient’s serum is largely unknown. Interestingly, the aHUS mutations often occur within the two most C-terminal domains of FH, which are the key regions for FH’s interactions with both C3b and the cellular surface (16, 17). This association between aHUS and mutations in FH CCPs 19 and 20 highlight s the importance of these regions for maintenance of complement regulation on host-cell surfaces.

We have generated an anti-FH agonistic antibody that is able to potentiate the effector functions of FH on human cells without affecting the bactericidal activity of the complement system (18). This potentiation is characterized by an improved affinity of FH for C3b and a reduced IC_50_ of C3b deposition and complement mediated lysis of sheep erythrocytes. The agonistic anti-FH was able to restore complement regulation in aHUS patient sera (18), but it remained unclear whether the observed effect was due to enhancement of both WT and mutant FH or if the anti-FH agonistic antibody potentiates only the WT FH in these heterozygous patient’s sera. Here, we have studied the effect of our potentiating anti-FH antibody on four naturally-occurring FH mutants that are associated with aHUS. Three of the four tested mutants were functionally restored to the normal level of regulation in the presence of the potentiating anti-FH. This demonstrates the agonistic activity of this antibody on WT and aHUS associated FH mutant proteins and provide *in vitro* proof of concept evidence for this agonistic antibody to be developed for the therapeutic treatment of aHUS caused by heterozygous mutations in FH.

## Material and Methods

### Antibodies, proteins and reagents

Recombinant human C-terminal HIS-tagged FH wild type (WT), or with mutations in CCP20; W1183L, V1197A, R1210C, G1194D, were provided by Gemini Therapeutics. The recombinantly produced FH mutants were over 95% pure as shown by SDS page (**Supplemental Fig. 1**).The recombinant expression of FH did not negatively affect the affinity of the potentiating anti-FH antibody (chimeric IgG4) to FH and was slightly better compared to the affinity of our antibody to plasma derived FH (pdFH) (18) and all recombinant FH had similar affinities (K_D_ ≈ < 1 nM, data not shown) for the potentiating antibody as determined by surface plasmon resonance (SPR).

pdFH, FB, FD and C3b were purchased from Complement Technology. In addition, for the C3b binding experiment in SPR, pdFH was obtained by in house isolation as previously described (19).

Chimeric versions (human IgG4) of anti-FH.07 (18) and anti-FH.07.1 have been provided by Gemini Therapeutics. Compared to previously published anti-FH.07 this paper makes use of anti-FH.07.1, further referred to as the “potentiating anti-FH”. This antibody has a similar binding epitope and comparable function, as shown by competition ELISA and C3b deposition ELISA, performed as described previously (18) (**supplemental Fig. 2**).

Mouse-anti-human monoclonal antibodies anti-C3-19 (20), anti-FH.16 (directed against FH CCP-16, non-competing control), anti-FH.09 (directed against FH CCP-6, inhibiting control) (18), anti-IL6-8 (IgG control) (18), anti-CD46.3 and anti-CD55.1 (21) were produced in house and labeled as indicated. Proteins or antibodies were biotinylated according to the manufacturer’s instructions using EZ-Link Sulfo-NHS-LC-Biotin, No-Weigh Format (Thermo Scientific). Fluorescent labeling of antibodies was done with DyLight 488 or DyLight 647 Amine-Reactive Dye (Thermo Scientific) according to manufacturer’s instructions. Fab’ fragments of the monoclonal antibodies have been provided by Gemini Therapeutics or are generated by pepsin cleavage of in house produced antibodies as described previously (18).

Normal pooled serum (NPS) was obtained from > 30 healthy donors with informed, written consent, in accordance with Dutch regulations. This study was approved by the Sanquin Ethical advisory board in accordance with the Declaration of Helsinky. Serum was obtained by allowing blood to coagulate for 1 h at room temperature (RT) and collecting the supernatant after centrifugation at 950 × g for 10 min, pooled serum was aliquoted and stored at -80°C. FH depleted serum was purchased from Complement Technology. Eculizumab (Alexion Pharmaceutical) was obtained by collecting surplus from used Soliris injection bottles. Sheep erythrocytes (ES) were from Håtunalab. High-performance enzyme-linked immunosorbent assay (ELISA) buffer (HPE), streptavidin conjugated with poly– horseradish peroxidase (strep-poly-HRP), were from Sanquin Reagents. Anti-CD59 (MEM-43, FITC) was ordered from Thermo Scientific.

### ELISA

Unless stated otherwise, all ELISA incubation steps were performed for 1 hour at RT on a shaker. After each incubation, plates were washed 5 times with PBS containing 0.02% Tween-20 (PBS; Sanquin Diagnostics) using an ELISA washer (Biotek, 405 LSRS). All ELISAs were developed with 100 µl substrate solution per well containing 0.11 M sodium acetate (Merck), 0.1 mg/mL 3,5,3’,5’-tetramethylbenzidine (TMB, Merck) and 0.003% (v/v) H_2_O_2_ (Merck) diluted in milliQ (Merck). Reactions were stopped with 100 µL 0.2M H_2_SO_4_ (Merck). Time that reactions were incubated varies per ELISA and are mentioned below. Absorbance was measured at OD450_nm_ using a Synergy 2 Multi-Mode plate reader (BioTek Instruments) and corrected for the absorbance at OD540_nm_. All ELISA steps were performed with a final volume of 100 μL per well.

### C3b deposition on LPS

C3b deposition assay with NPS was performed as described previously (18). C3b deposition assay with FH depleted serum was performed as followed. Polysorp 96-wells microtiter plates (Nunc) were coated with Salmonella typhosa LPS (40 μg/mL, Sigma-Aldrich) in PBS, O/N at RT. After washing NPS or FH depleted serum was incubated in Veronal buffer (VB; 3 mM barbital, 1.8 mM sodium barbital, 145 mM NaCl, pH 7.4) containing 0.05% (w/v) gelatin, 5 mM MgCl2, 10 mM EGTA and 0.1% (w/v) Tween-20 in the presence or absence of FHs and/or anti-FH potentiating mAb at indicated concentrations. C3b deposition was detected with biotinylated mAb anti-C3.19 (0.55 μg/mL in HPE) followed by incubation with 0.0001% (v/v) strep-poly-HRP in HPE for 30 minutes. The ELISA was further developed as described above.

### SDS page, coommassie

To analyze the recombinant FH variants, **0**.**3 µg** of sample was denaturized in 4x LDS sample buffer (Invitrogen) and incubated for 10 min at 70°C. Samples were loaded on 4-12% bis-tris gel (Life Tech). Gel was run at 200 V for ∼55 min in MOPS buffer (Novex). Gel was stained for 20 minutes using instant blue stain (Expedeon) and unstained in MilliQ for approximately 16 hours. Gel was photographed using a Chemidoc (Bio-rad).

### Surface Plasmon Resonance (SPR)

All surface plasmon resonance (SPR) experiments were performed using a Biacore T200 (GE Healthcare) with either a research grade Series S Sensor Chip Protein G (GE healthcare) or Series S Sensor Chip CM5 (GE healthcare). Unless stated otherwise, SPR experiments were performed at 25°C using a flow rate of 15 µL/min and in phosphate buffered saline (PBS, Fresenius Kabi), pH 7.4 with 0.1% (w/v) Tween-20 (Merck) (PBS-T). Data were collected at a rate of 10 Hz and all SPR data were analyzed using Scrubber (v20c, Biologic).

### Antibody affinity for recombinant FH in SPR

For the assessment of affinity of the potentiating antibody for FH mutants, at the beginning of each cycle, the potentiating antibody was captured on a Series S Sensor Chip Protein G at a concentration of 1 µg/mL during 60 seconds, corresponding with a ∼200 response units (RU) signal increase, leaving a second flow cell blank as reference. After a stabilization period of 300 seconds, a titration of the FH of interest (155 kDa) was injected, in random order in a two-fold dilution range starting at 500 or 250 nM. The complex was allowed to associate for 600 seconds and dissociate for 1500 seconds. After each FH injection, the Protein G chip was regenerated for 30 seconds with 100 mM Glycin-HCl (Merck) pH 1.5 at 30 µL/min and the process was repeated starting a new cycle with the capture of the potentiating antibody. Affinity constants ware determined using kinetic modelling.

### Coupling C3b sensor chip for SPR

For the assessment of the affinity of FH to C3b or the decay accelerating activity of FH in the presence of various antibodies (Fab’ fragments), purified C3b was immobilized via amine-coupling on a series S CM5 Sensor Chip using standard methods. In short, flow channels were activated for 7 minutes with a 1-to-1 mixture of 0.1 M N-hydroxysuccinimide (GE healthcare) and 0.1 M 3-(N,N-dimethylamino) propyl-N-ethylcarbonadiimide (GE healthcare) at a flow rate of 5 µL/min. The reference flow channel was blank immobilized, the other was immobilized with C3b, diluted in 10 mM sodium acetate (BioRad), pH 5.0, with target immobilization response of 2000 RU. When desired RU signals were reached, the surfaces were blocked with a 7 min. injection of 1 M ethanolamide, pH 8.0 (GE healthcare) to finish immobilization.

### Affinity of FH to C3b in SPR

To asses binding of FH to C3b, FH was injected in two-fold dilution range, starting at 10µM, 5µM (in house obtained pdFH) or 625 nM (recombinant) FH, in random order, with a flow rate of 10 µL/min, at 37° C, over both reference and C3b coupled flow channels of the C3b coupled sensor chip described above and allowed to associate and dissociate for 60 seconds each. After each cycle the flow channels were regenerated for 10 seconds with 2 M NaCl (Merck) at 10 µL/min. To assess the effect of various potentiating antibodies on the binding affinity of FH to C3b, FH was similarly titrated in buffer containing a surplus of potentiating antibody Fab’ fragments; either 10 µM or 1.25 µM (2 times highest FH concentration in respective experiment). The obtained signals were corrected for the MW of FH (155 kDa) and the FH-anti-FH Fab’ fragment complex (205 kDa). Affinity constant (K_D_) was determined by plotting the affinity curve at equilibrium of binding.

### Decay acceleration activity (DAA) assay in SPR

To assess DAA of FH in the absence or presence of one of the various potentiating antibodies, a C3b coupled sensor chip as described above was used. Running buffer in this experiment was 10 mM HEPES with 150 mM NaCl, 1 mM MgCl_2_ and 0.005% (w/v) Tween-20, pH 7.4. Regeneration buffer was 10 mM HEPES with 150 mM NaCl, 3.4 mM Ethylenediaminetetraacetic acid (EDTA) and 0.005% (w/v) Tween-20, pH 7.4. To form the C3 convertase C3bBb, FB and FD were simultaneously injected at a concentration of respectively 600 nM and 100 nM for 180 seconds, corresponding with a ∼250 RU increase. After a stabilization period of 120 seconds (natural decay of the convertase), FH (50 or 12.5 nM) with or without the potentiating antibodies (100 or 200 nM) Fab’ fragments were injected for 180 seconds. Finally, buffer was flowed over for another 300 seconds. After each cycle the flow channels were regenerated for 30 seconds with regeneration buffer at 30 µL/min. Runs were performed without or with FB and FD capture, respectively the “buffer” or “FB and FD” conditions. To obtain the final superimposed DAA signal, the signals of the buffer conditions were subtracted from the corresponding FB and FD conditions.

### Hemolytic assay

Hemolytic assay was performed with some adjustments for the use of FH depleted serum. Each component described below was 25% (v/v) of total assay volume. In a U-shaped assay plate containing a 2-mm diameter glass bead (merck), FHs were diluted in VB + 0.05 % gelatin (Merck) (VBG) with or without an excess of potentiating anti-FH and titrated in two-fold. Sheep erythrocytes (SE) were washed with PBS and resuspended in VB + 5.8% sucrose (VBS), to contain 1.05 x 10^8^ cells/mL in final assay concentration. NPS and FH depleted serum were diluted in VBG + 0.1 mM EDTA, to be 10% (v/v) in final assay conditions. To activate the alternative pathway, VBG + 5 mM MgCl_2_ + 10 mM EGTA in final concentration was added to the wells. VBG with 10 mM EDTA was used as blank. Samples were incubated at 37°C for 75 minutes while shaking at 450 RPM (Eppendorf thermomixer). Lysis was stopped by adding 100 μL VBG followed by centrifugation (2.5 minutes, 1800 RPM/471 RCF, 7°C). Hemolysis was measured as absorbance of the supernatants at 412 nm, corrected for background absorbance measured at 690 nm, and expressed as percentage of the 100% lysis control (ES incubated with 0.85‰ (w/v) Saponin). Graphs were fitted using a nonlinear fit, [inhibitor] versus response with four parameters.

### C3 deposition on HAP-1 cells

HAP-1 cells (22) (Haplogen Genomics, Wien, Austria) deficient for CD46, CD55, and CD59 (deficient HAP-1 cells) (21) were cultured in Iscove’s Modified Dulbecco’s Medium (IMDM; Lonza) supplemented with 10% (v/v) fetal calf serum (FCS; Sigma-Aldrich), 100 U/mL penicillin (Invitrogen) and 100 μg/mL streptavidin (IMDM++; Invitrogen) at 37 °C and 5% CO_2_. Cells were washed with PBS and detached using Accutase (Sigma-Aldrich). Surface expression was checked by staining with anti-CD46-3-DyLight 488 (FITC), anti-CD55-1-DyLight 647 (APC), and anti-CD59-FITC conjugated antibodies and analysis using flow cytometry.

Complement activation and detection was performed as followed. In a U-shaped assay plate containing a 2-mm diameter glass bead, 7 x 10^4^ HAP-1 cells were incubated for 1 h at 37 °C while shaking at 350 RPM with 25% (v/v) NPS or FH depleted serum diluted in VBG supplemented with 10 mM MgCl_2_ and 20 mM EGTA (VBG^MgEGTA^) in the presence of equimolar (20 µg/mL) blocking anti-C5 (Eculizumab) to prevent lysis. FHs and/or potentiating anti-FH were added to the cells prior to addition of the serum. After incubation, deficient HAP-1 cells were washed with PBS supplemented with 0.5% (w/v) bovine serum albumin (BSA, Sigma-Aldrich) before staining in PBS containing 0.1% BSA. C3 deposition was detected by incubation of the HAP-1 cells with 1 µg/mL DyLight 488-conjugated or DyLight 647-conjugated anti-C3-19 for 30 min at room temperature. After incubation cells were washed 3 times with PBS containing 0.5% BSA and subsequently fixed 1% (w/v) paraformaldehyde (PFA, Merck Millipore) in PBS. LSR Canto II flow cytometer (BD Biosciences) was used for measuring and data analysis was performed using FlowJo software V10 (Treestar, Ashland, OR). Gating strategies are shown in **Supplemental Fig. 3**.

### Statistical analysis

Analysis and statistical tests were performed using GraphPad Prism version 8.0.2 (GraphPad Software).

## Results

### Potentiation of affinity of FH for C3b by potentiating anti-FH

We previously showed that our potentiating antibody can restore complement regulation in aHUS patient samples (18). To investigate whether our antibody can potentiate both WT as well as mutant FH we used recombinant variants of FH, namely wild type (WT) and aHUS-associated mutants W1183L, V1197A, R1210, and G1194D (> 95% pure, **Supplemental Fig 1**). First, we studied the potentiating effect of anti-FH.07.1 on FH binding to C3b. **Fig. 1** shows that the addition of anti-FH.07.1 fab’ fragments to pdFH increases the affinity for C3b by ∼ 3-fold compared to pdFH injection alone, with K_D_s of 1.9 and 6.0 µM respectively (**Fig. 1A, 1B**). We observed a clear titration dependent increase in binding of pdFH to C3b, confirming previous study (18). All tested full length recombinant FH mutants showed slightly (< 2-fold) weaker binding to C3b compared to recombinant WT FH. Addition of the potentiating anti-FH Fab’ fragments increased affinity to C3b of all tested FH mutants by 1.8 - 2.4 fold (**Fig. 1C, 1D, Table I**). In conclusion, these experiments show that the studied mutations in FH only slightly impair the binding to C3b and that all tested mutants are enhanced in their affinity to C3b by anti-FH.07.1.

**Table I:** Estimated affinity of recombinant FH for C3b with and without addition of anti-FH.07.1 Fab’ fragments

| FH variant | $K_D$ ($M^{-1}$), FH only * | $K_D$ ($M^{-1}$), FH + pot anti-FH Fab' * | Fold change** | n |
| --- | --- | --- | --- | --- |
| WT FH | $3.0 \times 10^{-6} \pm 2.0$ | $1.7 \times 10^{-6} \pm 1.2$ | $1.8 \pm 0.1$ | 4 |
| W1183L | $4.6 \times 10^{-6} \pm 2.5$ | $2.1 \times 10^{-6} \pm 1.1$ | $2.3 \pm 0.0$ | 2 |
| V1197A | $5.0 \times 10^{-6} \pm 2.9$ | $2.2 \times 10^{-6} \pm 1.1$ | $2.2 \pm 0.2$ | 2 |
| R1210C | $4.9 \times 10^{-6} \pm 3.0$ | $2.1 \times 10^{-6} \pm 1.4$ | $2.4 \pm 0.1$ | 2 |
| G1194D | $4.7 \times 10^{-6} \pm 2.1$ | $2.4 \times 10^{-6} \pm 1.4$ | $2.1 \pm 0.3$ | 2 |

\* affinities are estimates as experimental setup could not reach the required higher concentrations
\*\* improved affinity of FH + anti-FH.07.1 Fab' fragments over FH alone

**Figure 1:**
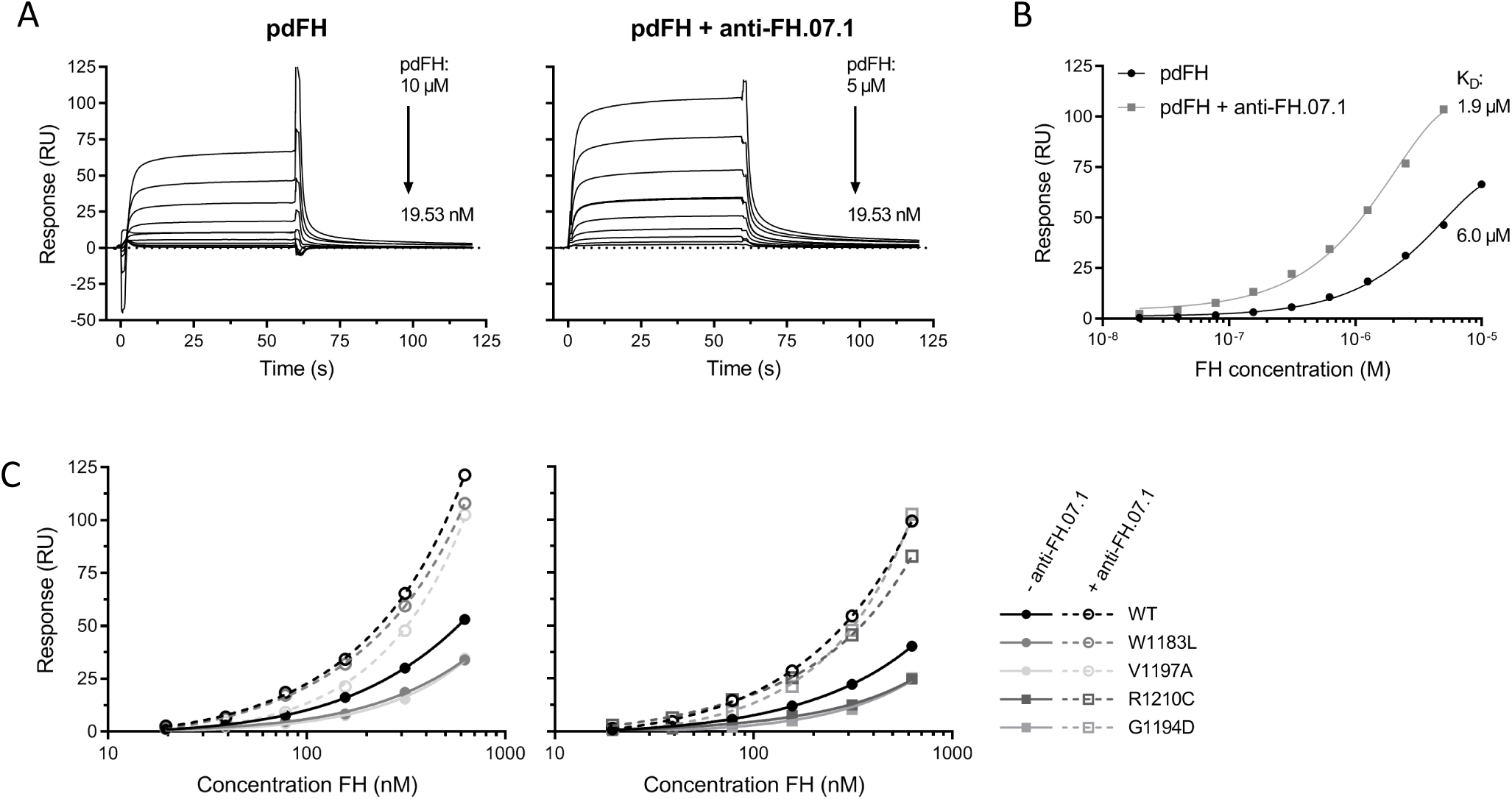
Affinity for C3b of mutant FHs is conserved and enhanced by potentiating antibody. The affinity of FH for C3b was measured using SPR. Measurements were performed on the Biacore T200 with CM5 chip coupled with ∼2000 RU C3b. (A) Sensorgrams of in house purified plasma derived FH (pdFH) binding to C3b. pdFH was titrated from 10 or 5 µM respectively, in 2 fold dilutions and flown over the chip without (left) or with (right) the presence of an excess (10 µM, at least 2 fold based on molar concentration) of anti-FH.07.1 Fab’ fragments. Sensorgrams were corrected for molecule size (155 KDa FH alone, 205 KDa FH + potentiating Fab’ fragment), and show an increased response upon addition of the anti-FH.07.1 Fab’ fragment. (B) Affinity curves, based on average response at equilibrium binding (ΔT 50-55 sec) in Fig. A, show an increase in binding upon addition of anti-FH.07.1 as presented by the estimated affinity K_D_ of 6.0 and 1.9 µM for pdFH alone or with addition of the anti-FH.07.1 Fab’ fragments respectively. (C) Affinity curves based on 2-fold titrations (625 – 39.0625 nM) of recombinant WT and mutant FH, corrected for molecule size, as described above, showing an increased response upon addition the anti-FH.07.1 Fab’ fragment (1.25 µM, dashed lines). Experiments are performed in two sets due to instrumental limitations. Figures are representative of n=2.

### All FH mutants show decay acceleration of AP convertase and are potentiated by potentiating anti-FH

Next, we studied the potential of the FH mutants to perform DAA of the AP convertase utilizing a SPR based assay (23). In this assay, we constructed the AP convertase C3bBb on the SPR chip by coupling C3b to the chip followed by incubation with FB and FD after which the AP convertases are formed (**Fig. 2A**, phase I). The AP convertases will naturally decay (**Fig. 2A**, phase II). As expected, this decay is increased upon injection of FH (**Fig. 2A**, phase III). With addition of anti-FH.07.1 Fab’ fragments pdFH showed increased DAA compared to pdFH alone as shown by the faster decrease in signal (**Fig. 2A**, grey dashed line). The anti-FH Fab’ fragments alone did not enhance DAA (**Fig. 2A**, black dashed line). Addition of FH binding Fab’ (anti-FH.16) or blocking Fab’ (anti-FH.09) (18) fragments results in respectively similar or less DAA compared to pdFH alone, confirming the potentiating effect of anti-FH.07.1 (**Fig. 2B**).

**Figure 2:**
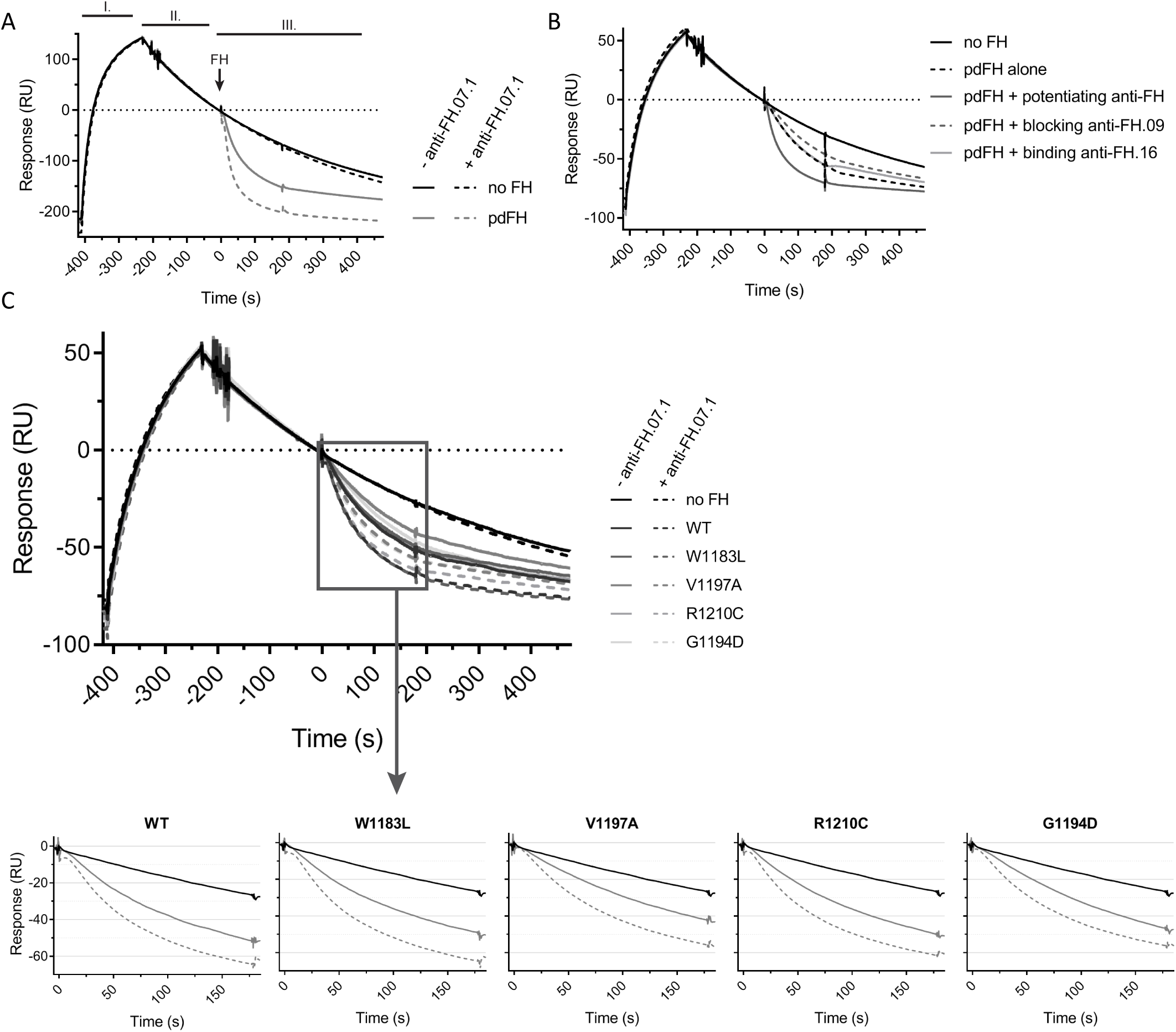
DAA of mutant FHs is enhanced by potentiating anti-FH. (A) In a SPR based setup FB and FD were flown over a sensor chip which was amine-coupled with ∼2000RU C3b to form C3bBb (convertase) complexes (phase I). Subsequently, a decline in signal indicated natural decay of the convertase as Bb is released from the coupled C3b (phase II). Injection of pdFH (50nM) causes accelerated decay (grey, solid line), as observed by a sudden further drop in response (phase III). The addition of the anti-FH.07.1 Fab’ fragment (200nM) to pdFH increases DAA of pdFH (grey, dashed line), whilst the Fab’ fragment alone does not affect the natural decay (black, dashed line). (B) With similar setup as described above, addition of anti-FH.07.1, FH blocking (anti-FH.09, (18)) or binding (anti-FH.16, (18)) anti-FH Fab’ fragments respectively increase, decrease or do not affect DAA of pdFH. (C) Injection of recombinant WT or FH mutants (12.5 nM) shows DAA. Addition of the anti-FH.07.1 Fab’ fragment (100nM) increases the DAA of all FHs. Enlargement of the DAA segment of SPR shows slight differences in DAA are observed between FH mutants. Addition of the anti-FH.07.1 Fab’ fragment (dashed lines) improves the DAA of all FH. Figures are average of duplicate runs and representative of n=8 (A), n=1 (B) or n=3 (C).

We then studied the potential of all tested FH mutants to accelerate the decay of the C3 convertase. All tested mutants did have DAA, although less efficient compared to WT FH (**Fig. 2C**, solid lines), and were all potentiated by addition of anti-FH.07.1 Fab’ fragments (**Fig. 2C**, dashed lines). Small differential effects of each tested FH mutant is observed. Without potentiation, WT FH, W1183L and R1210C show the highest activity in the DAA assay followed by, V1197A and G1194D, as determined by the final decrease in response. All tested FH mutants are similarly enhanced by the addition of anti-FH.07.1.

### The regulatory properties of most FH mutants can be enhanced by the potentiating anti-FH on cellular surfaces

In the SPR-based C3b binding and DAA assays, only small differences were detected comparing each of the four tested FH mutants with WT FH, so the potential to potentiate these mutants with anti-FH.07.1 on human cell surfaces was investigated. A cellular assay employing human HAP-1 cells was utilized in which all the membrane bound complement regulatory proteins (CD46, CD55 and CD59; these cells are naturally deficient of complement receptor 1) were knocked out (21). These cells are highly susceptible to complement activation when incubated with NPS, as shown by the high amount of C3 deposition (**Fig. 3a and Supplemental Fig. 3**). In this assay, addition of pdFH to the NPS decreases the C3 deposition, showing extra regulation from the added pdFH (**Fig. 3A and 3B**). Addition of the potentiating anti-FH to the NPS enhances the regulatory function of the present FH and as a consequence also decreases C3 deposition. These results indicate that HAP-1 cells deficient of all membrane bound complement regulatory proteins are highly dependent on FH binding to regulate complement deposition on their surface. The effects of the mutant FH proteins were investigated in this model both in isolation and in combination with the potentiating anti-FH. **Fig. 3C and 3D** shows that addition of the WT FH improved the regulatory balance on the deficient HAP-1 cells. None of the tested mutant FH variants was able to do so. Only a mild non-significant decrease in C3 deposition is observed compared to serum alone with addition of mutants V1197A, R1210C and G1194D (**Fig. 3C and 3D**, solid lines). Addition of anti-FH.07.1, which will bind to both serum FH from the NPS and the added recombinant FH, significantly increased regulation in all conditions (**Fig. 3C and 3D**, dashed lines).

**Figure 3:**
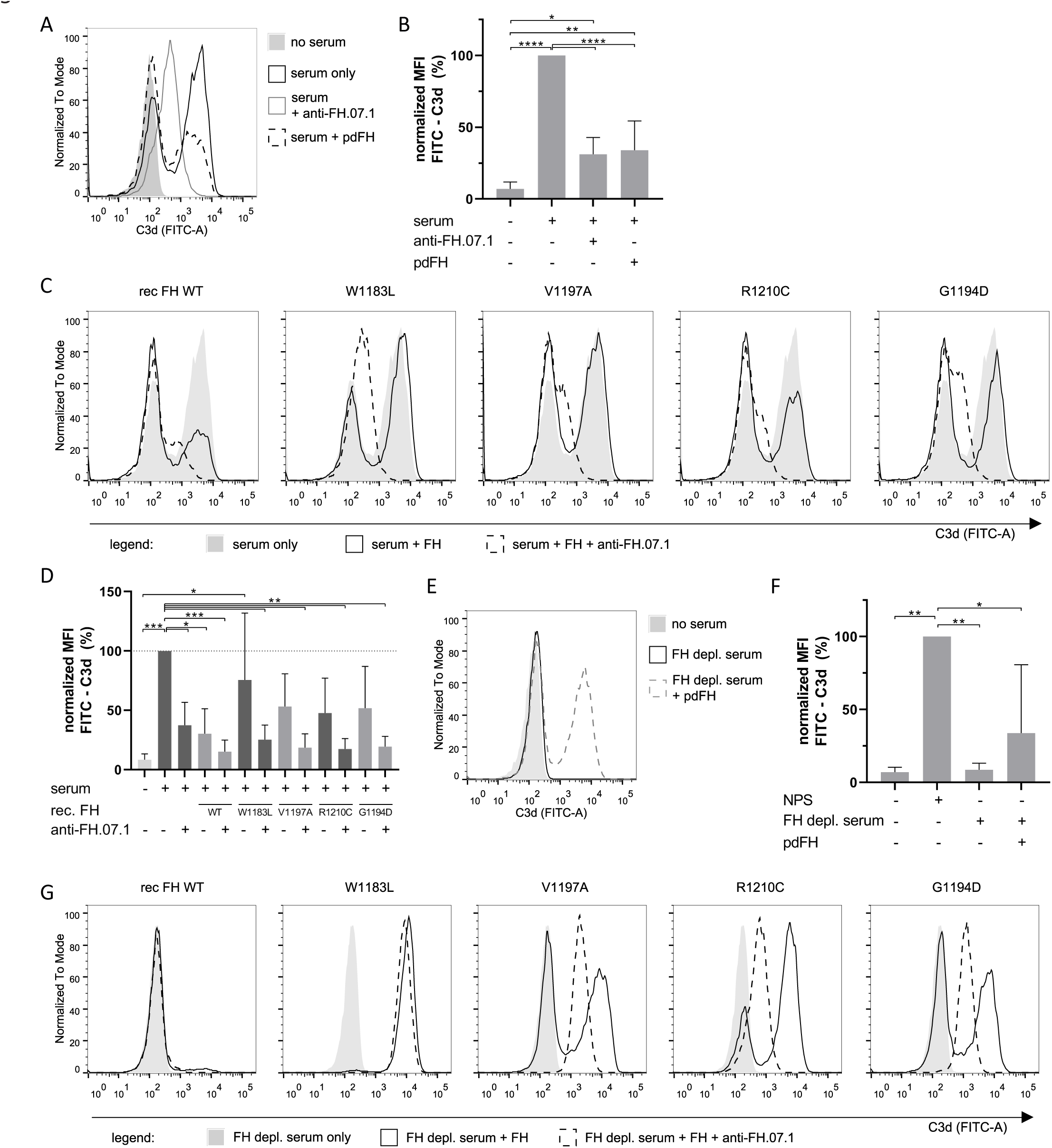
Deficient HAP-1 cells are protected by additional FH. (A) Flow cytometry analysis shows that HAP-1 cells deficient of all membrane bound complement regulators are sensitive to complement activation from NPS (25%) as shown by C3b deposition (black, solid line). Addition of the anti-FH.07.1 (75 µg/mL, grey line) or additional of pdFH (32.5 µg/mL, black, dashed line) shows control of C3 deposition. Histograms are a representative of n=4. (B) Normalized average MFI of conditions shown in Fig. A (n=4). (C) Addition of the recombinant WT or FH mutants (32.5 µg/mL, solid lines) show an increase in complement regulation on the HAP-1 cells deficient of membrane bound regulators as noted by a decreased C3 deposition compared to serum alone (grey, filled) and is further enhanced by the addition the anti-FH.07.1 (75 µg/mL, dashed lines). Histograms are a representative of n=4. (D) Normalized average MFI of conditions shown in Fig. C (n=4). (E) The addition of FH depleted serum (25%, black line) to deficient HAP-1 cells does not lead to C3b deposition as C3 is consumed in fluid phase in this serum before it can be deposited on the cells (25). Addition of pdFH (grey, dashed line) protects the fluid phase C3 consumption and thus renders the cells sensitive for complement activation as shown by increased C3b deposition. Histograms are a representative of n=3. (F) Normalized average MFI of conditions shown in Fig. E (n=3). (G) Addition of the recombinant WT or FH mutants (75 µg/mL, solid lines) to the FH depleted serum (25%) shows an increase in C3 deposition on the deficient HAP-1 compared to FH depleted serum alone (grey, filled), following the restoration of the regulation of fluid phase C3 consumption. Addition of the anti-FH.07.1 (75 µg/mL, dashed lines) next shows a decrease in C3 deposition compared to FH depleted serum and FH alone in flow cytometry. Histograms are a representative of n=3. Error bars represent standard deviation, * p < 0.1, ** p < 0.01, *** p < 0.001, analyzed by one-way ANOVA and Tukey’s multiple comparisons test.

To be able to fully distinguish the capacity of our antibodies to potentiate both WT as well as mutant FH, we further investigated complement activation on our HAP-1 cells deficient of complement regulators using FH depleted serum. We are unable to detect C3b deposition on the deficient HAP-1 cells when using FH depleted serum alone (**Fig. 3E and 3F**). This is consistent with data of others (12, 24, 25) and probably due to complete consumption of C3 in the FH depleted serum. Addition of physiological pdFH concentrations partly restores the C3 deposition to NPS control situation (**Fig. 3F**) as the fluid phase C3 conversion is also controlled. Addition of all FH mutants restores C3b deposition in this assay (**Fig. 3G**), indicating that all the tested mutant FH proteins restore the fluid phase regulation. **Fig. 3G** shows that FH V1197A, R1210C and G1194D restore C3b deposition equally well as pdFH, while addition of FH W1183L results in more C3b deposition. Addition of recombinant WT FH results in undetectable C3b deposition. We expect that addition of recombinant WT FH fully regulates fluid-phase C3 tick-over and inhibits complement deposition on the surface at the same time as we (**Supplemental Fig. 4**) and others (23) have observed that recombinant FH results in FH with better regulatory capacities than pdFH.

Addition of anti-FH.07.1, now acting only on the recombinant FH as no endogenous FH is present, shows that regulation by all mutant FH molecules, except FH W1183L, are enhanced as shown by decreased C3b deposition on the cell surface compared to the mutants alone (**Fig. 3G**, dashed lines). To conclude, V1197A, R1210C and G1194D are able to regulate C3 deposition on complement regulator deficient HAP-1 cells and are potentiated, by addition of anti-FH.07.1.

### C3b deposition is differentially controlled by all FH variants

To distinguish between the effects on fluid-phase and membrane regulation, we next investigated the impact of FH mutants on C3 regulation in serum, by assessing C3b deposition on LPS coated microtiter plates. To be able to solely asses the effect of our potentiating anti-FH on recombinant FH and not endogenous FH, this assay was performed using FH depleted serum. Similar to **Fig. 3E and 3F**, without FH in this assay we observe no C3 deposition because of low C3 levels caused by the natural consumption of C3 due to the tick-over of C3 into C3(H_2_O) (12, 25). Addition of all four tested FH mutants in increasing concentrations to the FH depleted serum first restores fluid phase complement regulation, resulting in measurable C3b deposition on LPS coated plates for all proteins up to ≤ 20 µg/mL supplemented FH (**Fig. 4A)**. This is followed by surface regulation upon further increasing FH concentration, resulting in a decreased C3b deposition.

**Figure 4:**
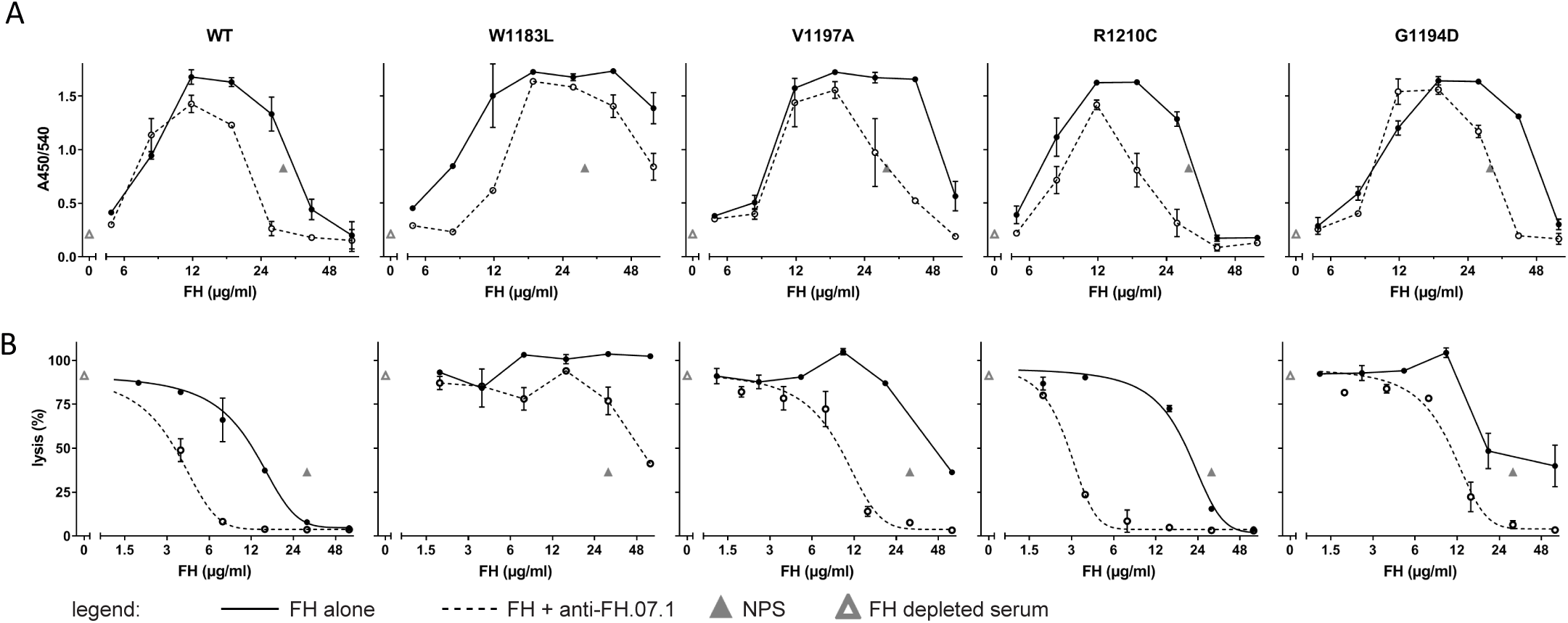
C3b deposition and hemolysis are differentially controlled by FH mutants. (A) LPS induced C3b deposition using FH depleted serum supplemented with either WT or FH mutants (solid lines) shows a concentration dependent increase in C3 deposition, indicating fluid phase regulation, followed by a decrease in C3b deposition, indicating surface regulation. Addition of the anti-FH.07.1 antibody (dashed lines) increases the regulatory activity of the most of the FH proteins. (B) Complement mediated lysis of SE incubated with FH depleted serum supplemented with either WT or FH mutants. Lysis is reduced by most of the FH proteins (solid lines). Addition of the anti-FH.07.1 antibody (dashed lines) improves the regulatory capacity of most of the FH proteins. The inhibition was fitted using a nonlinear fit, insufficient fit is not shown, instead the measured points are shown. Error bars represent standard deviation of experiments performed in duplicate and figures are representative of n=3.

Enhanced fluid phase regulation by any of the tested FH mutants was not observed in the presence of anti-FH07.1. However, there was a positive effect on surface regulation with the addition of anti-FH.07.1. Less FH is needed to restore the surface regulation of the C3b deposition, and thus the antibody enhances the FHs’ function (**Fig. 4A**, dashed lines). To conclude, V1197A, R1210C and G1194D are able to regulate C3 deposition on LPS in FH depleted serum alone and are potentiated, albeit differentially, by addition of the potentiating anti-FH.

### FH differentially regulate hemolysis in sheep erythrocyte hemolytic assay

To investigate if the FH mutations affect surface regulation of C3 deposition on cells similar as shown in an ELISA based assay, we employed the sheep erythrocyte (SE) hemolytic assay (26) with minor modifications to use FH depleted serum. Similar to our ELISA based assay, supplementation of recombinant WT FH restores complement regulation of FH depleted serum as shown by lysis of the SE. With addition of higher amounts of FH no more lysis is observed (**Fig. 4B**), indicating full protection. In this assay, the WT FH was most potent, with lowest amount of FH needed to restore protection. Compared to WT FH, the four mutant FHs required a higher FH concentration to inhibit lysis, indicating that the FH mutants are less potent in restoring complement regulation on the cell surface. Interestingly, anti-FH.07.1 increased the regulatory activity of all FH variants except W1183L, which was not able to regulate lysis even with addition of anti-FH.07.1. We only observe a reduction in lysis upon the highest, non-physiological, concentration. Overall, our data shows that anti-FH.07.1 is able to enhance the regulatory capacity of aHUS associated FH mutants V1197A, R1210Cand G1194D making this antibody an interesting drug candidate.

## Discussion

We had shown previously that our potentiating anti-FH antibody, anti-FH.07 was able to restore the FH activity in serum of aHUS patients (18). We currently investigated the effect of the anti-FH agonistic antibody, anti-FH.07.1 on four aHUS associated mutant variants of FH. However, in aHUS, patients are often heterozygous for FH mutations and it was unknown whether the antibody potentiated mutated FH in addition to WT FH. We now clarified that for three of the four tested mutants the anti-FH.07.1 monoclonal antibody increases the functionality of FH. This indicates that in aHUS patients carrying these select FH mutants, both the WT and the mutated FH protein will be potentiated.

Using SPR we confirmed that anti-FH.07.1 was binding to the recombinant FHs with comparable affinities. The affinities were around ten times higher than pdFH binding (data not shown). This increased affinity could be caused by the HIS-tag on the FH, which is known to cause artefactual enhanced binding on CM5 sensors, as used in this SPR setup (27). In addition, the tested FH mutants showed slight decrease in affinity for C3b compared to WT FH. Previous studies have shown conflicting results towards the effect of studied mutation on binding to C3b. Reports have shown variable effects of the W1183L, R1210C and V1197A mutations resulting in reduced C3b binding. These studies include the use of patient derived material, or recombinant constructs comprising CCPs 8-20 (FH8-20) or CCPs 18-20 (FH18-20) and the use of various assays (26, 28–31). Mutant G1194D has not been functionally studied before. The recombinant full length FH mutants used in this study are different than those used in previous studies and as a result, we cannot directly compare affinities. However, it seems that the involvement of CCPs 1-4, which contain an important C3 binding region (7, 11), in our full length FH dampens the effect on C3b binding as we see only a slight reduction in affinity (< 2-fold) between WT and four tested mutants which is in contrast to studies using FH8-20 or FH18-20 which see ≤ 60% reduction in affinity compared to the WT fragment (28–31). However, this does not explain the discrepancy between the results of Sanchez-Corral et. al and this study. Sanchez-Corral has isolated full length FH from healthy volunteers or patient material containing mutations W1183L, V1197A or R1210C (31). This studies indicates a ∼4-fold reduction in affinity towards C3b while we observe only moderate reduction. As described before, our recombinant proteins contain a HIS-tag which might have influenced our SPR based assay (27). However, as our WT FH also contains this HIS tag it might also be the sample preparation or other causes that affect the functionality of these mutants.

The addition of anti-FH.07.1 increases the affinity of all mutants for C3b with affinities that approach the affinity of WT FH for C3b. Anti-FH.07.1, which binds to CCP-18 (18), might alter the C3b binding interface of the C-terminal domains in such a way that CCP20 binding to the surface is less important. Possibly, the antibody affects the proposed “closed” or “folded” state of native FH (32, 33), allowing better binding to C3b. More research into the effect of our antibody on FH’s binding interaction with C3b and potential conformation changes are still needed to fully understand the mechanisms of action. Using the same C3b coupled SPR chip we observed very similar capacity for DAA of all FH mutants. As we have shown similar C3b affinity between mutants in our SPR based setup and co-factor activity relies on CCP domain 1-4 (7, 11) this data is in line with expectations. All four tested FH mutants are equally enhanced and only very small effects of the mutations are noticed.

The effects of the mutations are more pronounced in the experiments involving serum. Studies show that aHUS patients carrying the specific mutations used in this study (W1183L, R1210C, V1197A) do not present with consumption of complement (hypocomplementemia) (26, 30, 31). As mentioned above, it is known that CCPs 1-4 of factor H are responsible for co-factor activity and fluid phase regulation (7) and these domains are not affected in the these FH mutants (26). Our study shows similar results, all our mutants are able to control fluid phase regulation. However, especially W1183L seems to be able to control of fluid phase regulation better than other FH mutants and even WT FH.

The results of complement regulation on the surface using an ELISA format were in agreement with the results of the hemolytic assay. Mutant W1183L was only able to regulate some C3 deposition upon FH potentiation, when added in supra-physiological concentration. R1210C performed in these assays equally well as WT FH did, while this mutant is known to have reduced affinity for C3b and cellular surfaces (28).The results were unexpected, because this mutation has been repeatedly found in aHUS, C3G, as well as AMD patients (34). An explanation could be that in patient material it was found that the introduced cysteine residue forms disulfide bond between FH and albumin, forming a 210 KDa band in serum isolated FH (31). This phenotype can explain the cause of disease in patients, as the FH-HSA is functionally impaired and affects the ability of the patient serum to protect against SE lysis (31), and similar protein interactions with other proteins have not been identified to occur during the production of the recombinant mutant protein.

For all mutants, except G1194D, it has been shown that CCP8-20 fragments are affected in their capacity to bind to heparin or human umbilical vein endothelial cells (HUVEC) (28, 30), as well as the reduced ability of patient serum or recombinant FH fragments to respectively protect against or inhibit SE lysis (31, 35). This is in line with our own observations, except for R1210C, which performed equal to WT FH in our hemolytic assay. W1183L seems much less potent in controlling complement regulation on surfaces compared to the other mutants. Considering the function of studied mutations, other studies are not always in agreement. While one structural study of the FH18-20 fragment shows that the W1183 domain in CCP20 is directly involved in C3b/d binding (29), other studies demonstrate the close involvement of W1183 in binding to sialic acid (36). Mutations R1210C, V1197A and G1194D result in destabilization of the tertiary structure (29), while V1197 is also shown to be important for binding to sialic acid (36). However, it is also noted that other ligands might have differential interactions in this binding region, such as heparin sulphates compared to sialic acid (36, 37) and is explained by the various effector functions of FH on different cells types, such as endothelial cells, glomerular membrane, retinal pigment epithelium, Bruch’s membrane in the eye, platelets or erythrocytes (24, 37, 38). A dual interaction of FH to C3b and sialic acid residues is crucial for protection of cells against unwanted complement activation on human cell surfaces (38). W1183 is involved in both processes and it has been shown that in competition assays, the patient-associated W1183L aHUS mutation mostly affects the ability to compete with WT FH (38, 39). In these studies, Hyvärinen et al. show that removal of sialic acid from HUVECs and platelets showed minimal effect on the competition of this FH mutant FH18-20 compared to WT FH18-20 with WT FH, showing that the sialic acid was crucial for this interaction (38).

In conclusion, we have shown that four select recombinantly expressed aHUS associated FH mutant proteins can be enhanced in their binding to C3b and decay accelerating activity by our potentiating antibody. In addition, this anti-FH antibody can potentiate three of the four FH mutants in their regulatory function on the cellular surface. Taken together, this research confirms that our antibody can potentially be used to enhance the FH of aHUS patients who carry these mutations.

## Acknowledgments

We thank Arnoud Marquart, Richard Pouw and Christoph Schmidt for their kind help with methods and the analysis of data. We thank Dr. Erfan Nur for his kind help obtaining Eculizumab.

## Figure legends

**Supplemental Figure 1: Recombinantly produced FHs are pure**

Expression and isolation of FH resulted in pure, full-length FH. No contamination or breakdown products are detected in the recombinant FH fractions as shown by SDS page with coomassie blue staining.

**Supplemental Figure 2: The anti-FH.07.1 is comparable to anti-FH.07**

(A) Anti-FH.07 (18) and anti.FH.07.1 (potentiating anti-FH, this research) share the same binding epitope on FH, as shown by competition ELISA. Either anti-FH.07, anti-FH.07.1, or a non-competing anti-FH (anti-FH.16, (18)) is coated as a capture of biotinylated FH. All abovementioned anti-FH antibodies are also included as a competitor for binding of biotinylated FH, plus a non-competing control isotype control (IgG control) (B) The anti-FH.07.1 is comparable in function to anti-FH.07 as shown by the inhibition of LPS activated C3b deposition in NPS. Error bars represent standard deviation of experiments performed in duplicate and figures are representative of n=2 and n=4 for respectively Fig. A and Fig B.

**Supplemental Figure 3: Gating strategy to determine level of C3b deposition on deficient HAP-1 cells**

C3 deposition on HAP-1 cells deficient of all membrane bound complement regulators (Hap-1 KO CD46/CD55/CD59, (21)) incubated without (top panel) or with NPS (25%, bottom panel) was analyzed by flow cytometry using FITC or APC labeled anti-C3d (C3-19). Gating strategy was as followed: Deficient HAP-1 cells were gated based on size and granularity using FSC-A vs SSC-A to eliminate debris and clumped cells (plot 1). Single cells were sub-gated using SSC-H and SSC-W (plot 2) and subsequent FSC-H and FSC-W (plot 3), Continuity of measurement was checked by plotting the acquisition time (x-axis) to the APC-A signal, “time-gate” (plot 4), Cells positive for C3b were visualized using normalized histograms of either FITC-A or APC-A, “C3b deposition” (plot 5). SSC-A: side scatter area, FSC-A: forward scatter area, FSC-H: forward scatter height, FSC-W: forward scatter width, SSC-H: side scatter height, SSC-W: side scatter width.

**Supplemental Figure 4: Recombinant WT FH has improved protective function in hemolytic assay compared to pdFH**

Recombinant WT FH has a higher regulatory potential as shown by complement mediated lysis of SE incubated with FH depleted serum supplemented with either pdFH or recombinant WT FH. Lysis is reduced by addition of FH, less recombinant WT FH is needed to reach inhibit lysis compared to pdFH. The inhibition was fitted using a nonlinear fit. Error bars represent standard deviation, experiments were performed in duplicate and figures are representative of n=3.

