## Supplemental figures 1-4 for "Unravelling the effect of a potentiating anti-Factor H antibody on atypical hemolytic uremic syndrome associated factor H variants"

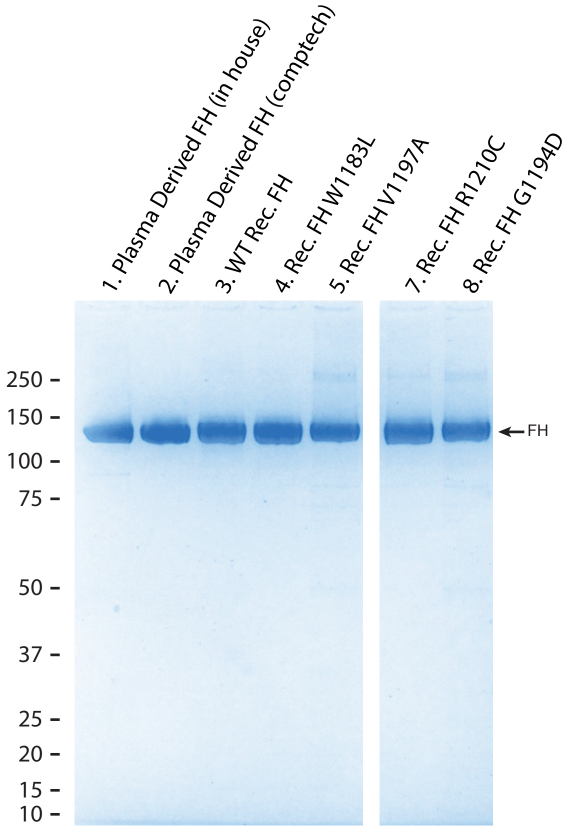


**Supplemental Figure 1: Recombinantly produced FHs are pure**

Expression and isolation of FH resulted in pure, full-length FH. No contamination or breakdown products are detected in the recombinant FH fractions as shown by SDS page with coomassie blue staining.

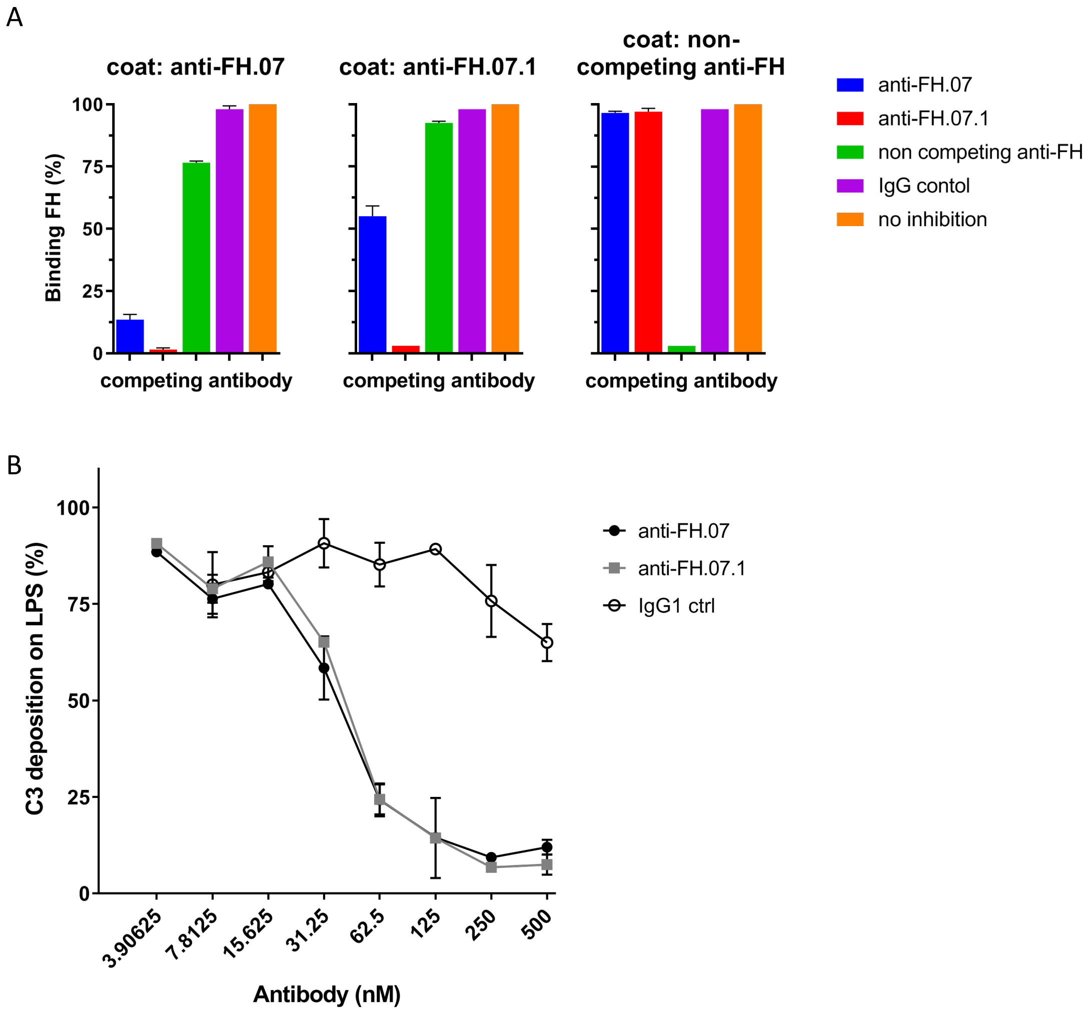


**Supplemental Figure 2: The anti-FH.07.1 is comparable to anti-FH.07**

(A) Anti-FH.07 (18) and anti.FH.07.1 (potentiating anti-FH, this research) share the same binding epitope on FH, as shown by competition ELISA. Either anti-FH.07, anti-FH.07.1, or a non-competing anti-FH (anti-FH.16, (18)) is coated as a capture of biotinylated FH. All abovementioned anti-FH antibodies are also included as a competitor for binding of biotinylated FH, plus a non-competing control isotype control (IgG control) (B) The novel potentiating anti-FH is comparable in function to anti-FH.07 as shown by the inhibition of LPS activated C3b deposition in NPS. Error bars represent standard deviation of experiments performed in duplicate and figures are representative of n=2 and n=4 for respectively Fig. A and Fig B.

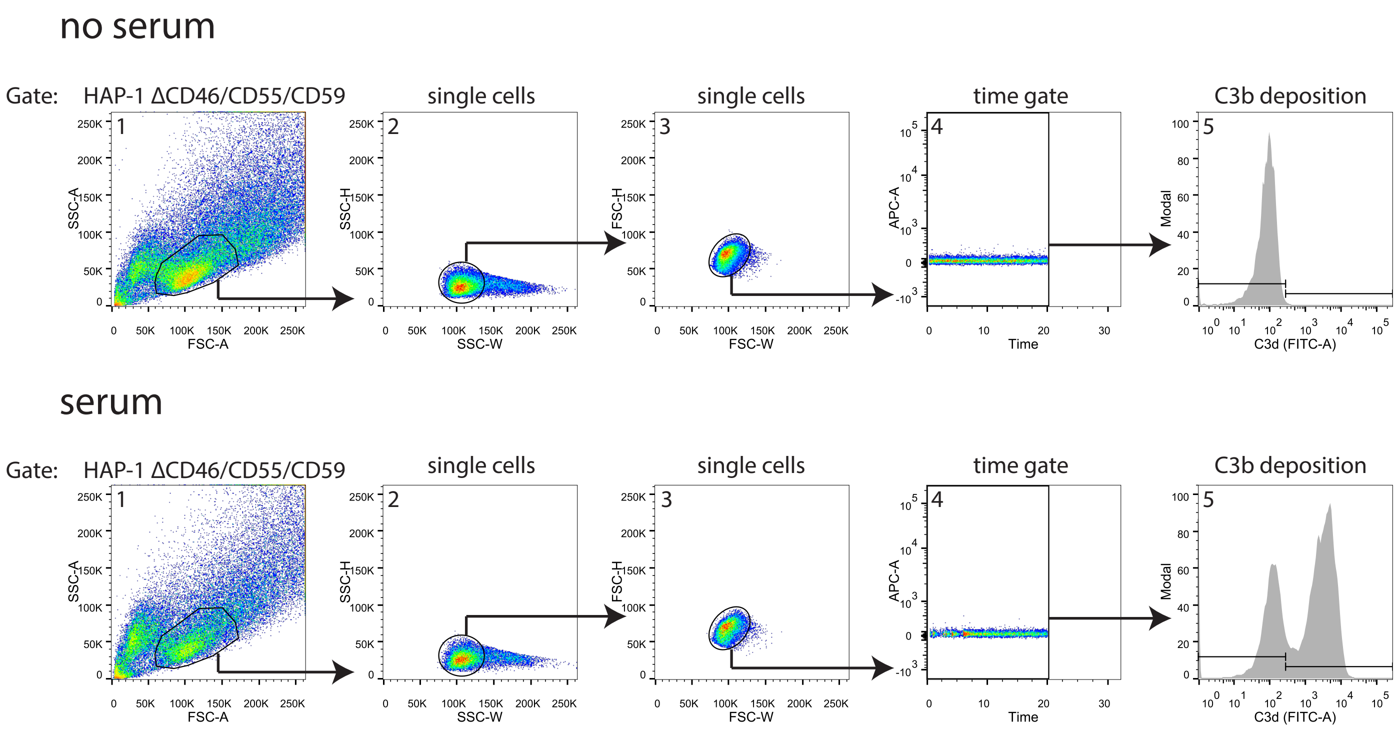


**Supplemental Figure 3: Gating strategy to determine level of C3b deposition on deficient HAP-1 cells**

C3 deposition on HAP-1 cells deficient of all membrane bound complement regulators (Hap-1 KO CD46/CD55/CD59, (21)) incubated without (top panel) or with NPS (25%, bottom panel) was analyzed by flow cytometry using FITC or APC labeled anti-C3d (C3-19). Gating strategy was as followed: Deficient HAP-1 cells were gated based on size and granularity using FSC-A vs SSC-A to eliminate debris and clumped cells (plot 1). Single cells were sub-gated using SSC-H and SSC-W (plot 2) and subsequent FSC-H and FSC-W (plot 3), Continuity of measurement was checked by plotting the acquisition time (x-axis) to the APC-A signal, "time-gate" (plot 4), Cells positive for C3b were visualized using normalized histograms of either FITC-A or APC-A, "C3b deposition" (plot 5). SSC-A: side scatter area, FSC-A: forward scatter area, FSC-H: forward scatter height, FSC-W: forward scatter width, SSC-H: side scatter height, SSC-W: side scatter width.

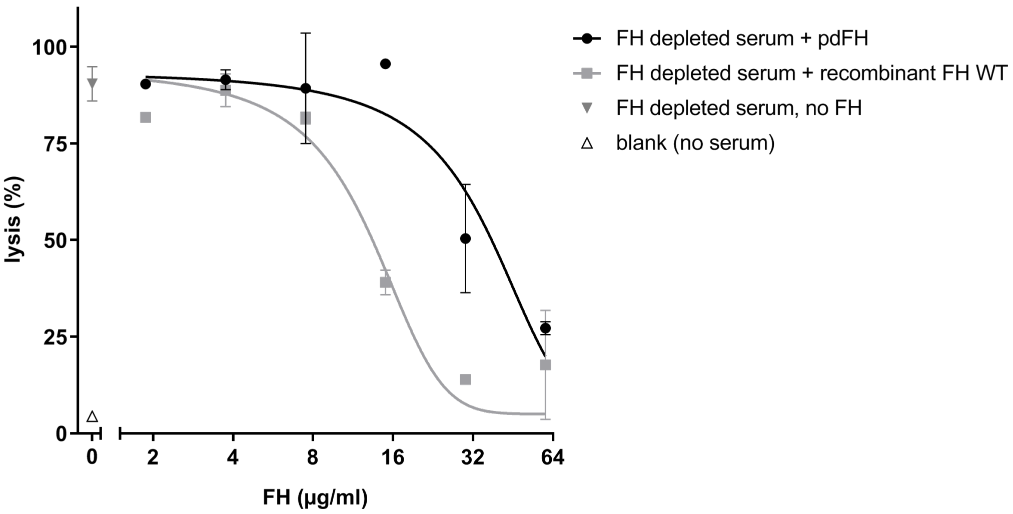


**Supplemental Figure 4: Recombinant WT FH has improved protective function in hemolytic assay compared to pdFH**

Recombinant WT FH has a higher regulatory potential as shown by complement mediated lysis of SE incubated with FH depleted serum supplemented with either pdFH or recombinant WT FH. Lysis is reduced by addition of FH, less recombinant WT FH is needed to reach inhibit lysis compared to pdFH. The inhibition was fitted using a nonlinear fit. Error bars represent standard deviation, experiments were performed in duplicate and figures are representative of n=3.
